# The hollow truth: ethylene-triggered ROS, PCD, senescence, and autophagy drive hollow stem formation

**DOI:** 10.1101/2025.03.06.641815

**Authors:** Mengxiao Yan, Weijuan Fan, Wei Yang, Jiamin Zhao, Yinghui Meng, Wuyu Zhou, Haiyan Zhuang, Ziyin Xu, Yuqin Wang, Qingjun Huang, Ling Yuan, Hongxia Wang, Jun Yang

**Author notes:** These authors contributed equally to this work (Mengxiao Yan, Weijuan Fan). Correspondence: H.X.W. and J.Y.

## Abstract

Hollow stems have independently evolved multiple times across the plant kingdom, and play crucial roles in plant development and various environmental adaptations. However, the mechanisms underlying stem hollowness remain poorly understood. Water spinach (*Ipomoea aquatica*) is one of the few hollow-stemmed plants in the *Convolvulaceae* family, and its hollow stems are essential for thriving in aquatic environments. Using histochemical staining and transcriptome analysis, we identified programmed cell death (PCD) is involved in cavity formation of water spinach shoot tips. Single-cell transcriptome analysis further revealed that ethylene and reactive oxygen species (ROS) drive and regulate this process by activating transcription factors *IaNAC074*, *IaNAC087*, *IaNAC029*, *IaNTL9*, and *IaTGA9*, which initiate PCD, senescence, and autophagy, collectively controlling cell death. These findings were validated through treatments with ethylene and ROS reagents in water spinach and transient expression in tobacco. Additionally, transcriptomic data suggest that these mechanisms may also play a role in hollow stem formation in bamboo, highlighting the conservation of PCD regulatory mechanisms in plant development. This work not only fills a major knowledge gap in the adaptive mechanisms of hollow stem formation but also opens new avenues for agricultural and ecological applications, offering strategies to enhance crop tolerance to flooding and accelerate crop growth.

## Introduction

Interestingly, hollow stems are a common trait across the plant kingdom, originating independently multiple times in various taxonomic groups. This trait can be observed in a wide range of plant groups, including some *Equisetum* species among ferns, bamboo and rice (Poaceae), *Eleocharis dulcis* (Cyperaceae), *Alisma canaliculatum* (Alismataceae) among monocots, and some *Ranunculus* species (Ranunculaceae), *Rorippa islandica* (Brassicaceae), *Alternanthera philoxeroides* (Amaranthaceae), *Ipomoea aquatica* (Convolvulaceae), *Artemisia leucophila* (Asteraceae) among eudicots ^1–3^. Usually, the parenchyma cells of the stem pith undergo death (a process known as pith autolysis or pithiness) in these plants, resulting in the formation of air-filled cavities or hollow stems (Carr et al. 1995). The hollow structure of plant stems plays a significant role in growth, development, and environmental adaptation. Since stem pith autolysis facilitates rapid growth while providing sufficient structural support, such as in bamboos ^4^, and acts as carbon sinks for growth demand ^5^. Stem pith autolysis also responses to fluctuations in various environmental factors. It has been reported that drought ^6^, waterlogging ^7^, water flow ^8^, low light ^5^, temperature ^2^, and nutrient deficiency ^9^ induce stem pith autolysis. This widespread occurrence highlights the evolutionary advantage and adaptability that hollow stems confer to plants in differing ecological niches. However, in agriculture, pith autolysis is often an undesirable trait as it can lead to decreased quality and reduced crop yields. This issue is commonly seen in the stems of broccoli and sweet sorghum, the roots of radish and turnip, and the fruits of cucumber and pear ^10–14^. Therefore, investigating the underlying mechanisms of pith autolysis is not only deepens our understanding of adaptive evolution in plants but also holds potential for addresses agricultural production challenges.

Water spinach (*Ipomoea aquatica* Forsk.), a commonly grown leafy vegetable, is a rich source of vitamins, minerals, proteins, fibers, carotenes and flavonoids with many health benefits ^15^. It is only true aquatic plant in genus *Ipomoea*, with roots on mud and floating stems on water (Fig. 1a) ^16^. Water spinach has characteristic hollow stem, which is an important trait to adapt the aquatic environment. The hollow stem can allow the shoot to float on the water surface, thus ensuring the plant can perform photosynthesis and gas exchange. At the same time, hollow stem can also ensure gas exchange between the plants on the water surface and the roots underwater, addressing the issue of low oxygen stress for the roots ^17^. Among the species within genus *Ipomoea*, the majority possess solid stems ^16,18^—a feature commonly associated with terrestrial growth habits. The hollow stem of water spinach is a notable exception. This evolutionary trait, presumed to have independently evolved from ancestors with solid stems, represents water spinach’s ingenious adaptation to aquatic environment. Water spinach could serve as an ideal model for studying the formation of hollow stems in dicotyledonous plants, providing insights into the plant adaptive evolution from terrestrial to aquatic environments in the meantime. In addition, water spinach is a close relative of sweetpotato (*Ipomoea batatas* (L.) Lam.), a staple crop with nutritious storage roots ^16^. Knowledge on pith autolysis in water spinach also holds significant importance for understanding and addressing the issue of pithiness in the storage roots of sweetpotato during senesces and storage ^19^.

**Fig 1.**
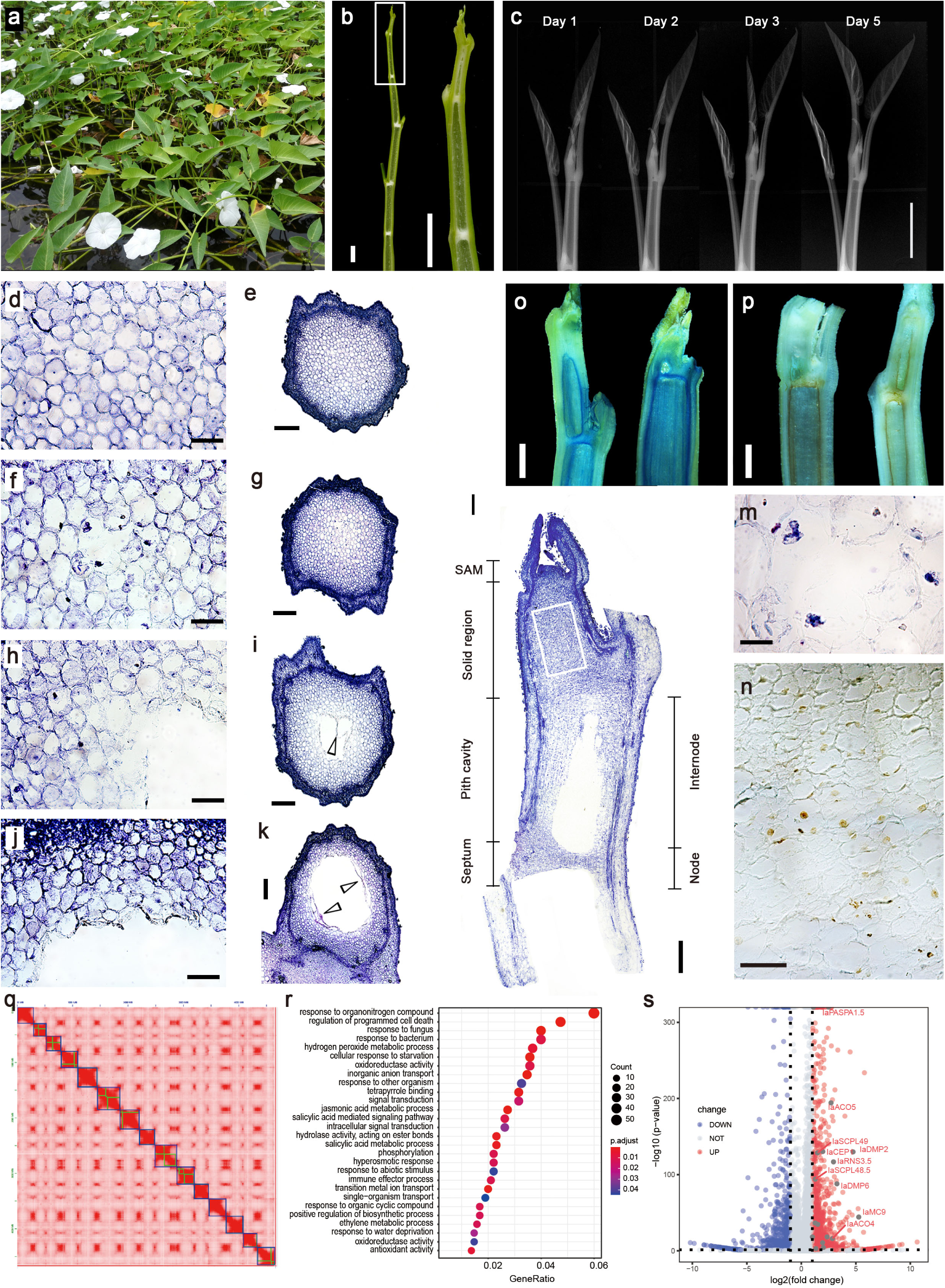
Anatomical characterization and transcriptome analysis of pith cavity development in water spinach. a. Water spinach floats on the water surface due to its hollow stems, enabling photosynthesis and gas exchange; b. Longitudinal section of the water spinach shoot, showing hollow internodes and solid septums at the nodes; the right panel is a magnified view of the boxed area in the left panel. Scale bars, 5 mm. c. X-ray scanning displaying the process of cavity development. Scale bar, 5 mm. d-l, Paraffin sections of stem tips stained with toluidine blue: d-e, Cross-sections of the solid tip showing intact cell walls and nuclei. Scale bars, 50 μm (d) and 200 μm (e). f-g, Cross-sections at the initiation of cavity formation, showing central pith cells with broken cell walls and gradually rupturing nuclei. Scale bars, 50 μm (f) and 200 μm (g). h-i, Cross-sections during cavity development, where outer cavity cells die, cell walls degrade and detach (indicated by arrows), and nuclei dissolved resulting in the absence of nuclei. Scale bars, 50 μm (h) and 200 μm (i). j-k, Cross-sections at the mature stage of cavity development, with multiple layers of detached dead cells (indicated by arrows) with degraded cell walls and absent nuclei. Scale bars, 50 μm (j) and 200 μm (k). l, Longitudinal section of the shoot tip, showing the solid apical meristem and cavity development at the base of the first internode, with solid septum at the nodes and hollow internodes. m, Magnified view of panel (f) showing the process of nuclear shrinkage and fragmentation. Scale bar, 20 μm. n, TUNEL staining concentrated in the nuclei of cells between the meristem and the cavity, as shown in the area outlined by the white box in panel l. Scale bar, 50 μm. o, Trypan blue staining localized around the cavity region of the shoot tip. Scale bar, 1 mm. p, DAB staining concentrated around the cavity region of the shoot tip. Scale bar, 1 mm. q, Hi-C heatmap of the newly assembled genome, displaying 15 chromosomes. r, GO term enrichment of upregulated genes in the cavity region relative to the solid region. s, Genes involved in PCD and ethylene biosynthesis are upregulated in the cavity region of the shoot tips.

As of now, research on the mechanisms of stem hollowing in plants remains scarce, with only a single study reported on hollow stem formation in bamboo ^20^. In herbaceous plants, the hollow stem is thought to from the autolysis of a plant’s storage pith ^5,21^. Pith autolysis was reported in many plants, including aerenchyma formation in stems of aquatic and wetland plants ^1,22^ and crop roots ^23–25^, and root pithiness in radish ^12^. Pith autolysis is potentially regulated by ethylene, calcium, and reactive oxygen species (ROS)-mediated signaling pathway, and programmed cell death (PCD)-executing enzymes are the most common downstream causes of pithiness-like phenotypes ^20,26,27^. In recent years, several transcription factors (TFs) have been found to act as master regulators of pithiness or PCD executing. In sorghum, *SbNAC074* induces programmed death of stem pith parenchyma cells through cell autolytic enzymes ^11^. *ANAC074* also promotes cell death in the stigma of Arabidopsis ^28^. In radish root, the elevated oxidative stress or hypoxia that activates *RsNAC013* for mitochondrial signaling, leading to cell death of the xylem parenchyma during the root-thickening process ^12^. In Arabidopsis, *ANAC033*, *ANAC046*, *ANAC087* and *ZAT14* mediate PCD in the root cap ^29–31^. However, due to the lack of research, the mechanisms underlying the formation of hollow stems in plants remain unclear.

To investigate the mechanism behind hollow stem formation in water spinach, we employed a comprehensive approach involving morphological anatomy, single-cell transcriptomics, and molecular validation. Histochemical staining and transcriptome analysis between hollow and solid stems demonstrated that PCD is central to pith cavity formation. Further single-cell transcriptomics revealed that ethylene-triggered ROS, along with PCD, senescence, and autophagy, collectively drive hollow stem formation. These mechanism also play a role in hollow stem formation in bamboo, indicating that the ethylene-ROS-PCD regulatory network may be conserved across plant species. Hollow stems can not only enhance crop tolerance to flooding, but also accelerate growth, improve resource use efficiency, and ultimately increase yield. Therefore, this study may inspire applications of this mechanism in other crops to enhance adaptability to environmental changes.

## Results

### Morphological and anatomical characterization of pith cavity development

The morphological structure of the stems of water spinach is similar with that of bamboo, characterized by hollow internodes interspersed with solid nodes (Fig. 1b). In water spinach, the top of shoot tip is solid, comprising meristem and pith (Fig. 1e, l). The cavity developed in the pith beneath meristem (Fig. 1l). Continuous X-ray scans have revealed that initially, the first internode of shoot tip is solid. Over time, a cavity begins to develop at the base of the first internode, gradually enlarging until a complete node and internode are formed (Fig. 1c).

### PCD plays an important role in the pith cavity formation

To investigate the fine structure of the cavity region and the mechanisms underlying cavity formation, we performed histochemical staining and observation on cross and longitudinal sections of shoot tips. In the solid regions of shoot tips, the cell nuclei were round, with intact nuclear membranes, and the nucleoli centrally located within the nuclei (Fig. 1d). During the early stages of cavity development, the broken cell walls, and nuclear shrinkage and fragmentation were observed in central pith cells (Fig. 1f, m). As the cavity development progressed, the nuclei dissolved, and the cell walls broke down and detached (Fig. 1h-k), indicating that the cells in the cavity regions had undergone cell death. In contrast, the cells surrounding the cavity retained intact nuclei and cell walls (Fig. 1f, h, j). These features suggested that the formation of the cavity is associated with cell death.

Therefore, we employed further histochemical staining and immunostaining techniques to verify whether programmed cell death occurs during cavity development. Since trypan blue is a colorimetric dye that stains dead cells with a blue color ^32^, we dropped trypan blue solution to cover the surface of longitudinally bisected shoot tips. The cutting surface appeared light blue as wound induced cell death (Fig. 1o). While the inner walls surrounding the cavity showed a deeper blue compared to the cutting surface, indicating that cell death has occurred around pith cavity (Fig. 1o). Since ROS are important regulatory signals in programmed cell death ^33^, we performed 3,3-diaminobenzidine (DAB) staining to detect for ROS accumulation (H_2_O_2_) ^34^. In the shoot tip of water spinach, DAB-stained cells were found around the cavity regions (Fig. 1p). This result indicated that ROS accumulated during cavity development. We also performed terminal deoxynucleotidyl transferase-mediated dUTP nick end labeling (TUNEL) staining, which is a method for staining cells that have undergone PCD by in situ labeling of DNA breaks in nuclei. The reaction is highly specific, only nuclei in apoptotic cells will be stained ^35^. Our observations showed that TUNEL staining was concentrated in the nuclei of cells located in the upper region of the cavity, indicating that these cells were undergoing programmed cell death (Fig. 1n). In contrast, the nuclei of cells in the cavity region lack TUNEL staining, suggesting that these cells had already undergone cell death (Fig. 1n).

To verify whether PCD is involved in cavity formation, we conducted a differential transcriptomic differential analysis between the solid area and cavity region of shoot tips (Supplementary Fig. 1a). Firstly, we assembled a high-quality genome of water spinach, resulting in the assembly of 15 chromosomes (Fig. 1q), to ensure comprehensive gene information for accurate transcriptome analysis. The BUSCO assessment indicated a completeness score of 96.5% (Supplementary Table 1), reflecting the accuracy and reliability of the assembled genome. Then, we conducted transcriptome analysis using this newly assembled genome as a reference. A total of 1,175 differentially upregulated genes (padj < 0.05 & log2FoldChange > 1) in cavity regions were identified compared with solid region (sampled regions illustrated in Supplementary Fig. 1a). Gene Ontology (GO) enrichment analysis of these upregulated genes revealed significant enrichment of terms related to regulation of programmed cell death, hydrogen peroxide metabolic process, oxidoreductase activity, and ethylene metabolic process, etc. (Fig. 1r). Many genes involved in ethylene biosynthesis [1-aminocyclopropane-1-carboxylic acid (ACC) synthase (*IaACS*), *IaACO*; ACC oxidase (*IaACO*), *IaACS*], ROS production (respiratory burst oxidase homolog D, *IaRBOHD*), and PCD execution, such as protease (cysteine endopeptidase 1, *IaCEP1*; serine carboxypeptidase-like protein 48, *IaSCPL48*; serine carboxypeptidase-like protein 49, *IaSCPL49*; *aspartic proteinase A1*, *IaPASPA1*, metacaspase 3, *IaMC3*; metacaspase 9, *IaMC9*, gamma vacuolar processing enzyme, *IaγVPE*), and nuclease (*ribonuclease 3, IaRNS3*; *bifunctional nuclease*, *IaBFN1*; *endonuclease 2 IaENDO2*) were detected up regulated in hollow regions (Fig. 1s, Supplementary Table 2). Sweetpotato is a wild relative of water spinach with solid stems (Supplementary Fig. 1a). A differential transcriptome analysis between their shoot tips revealed 19,311 genes upregulated in water spinach (sampled regions illustrated in Supplementary Fig. 1a). GO enrichment analysis showed that these upregulated genes were significantly enriched in regulation of programmed cell death and oxidoreductase activity (Supplementary Fig. 1b). Therefore, both intraspecific and interspecific transcriptome analyses confirmed PCD is likely involved in and regulate the formation of hollow stems.

### Cell atlas of shoot tip from water spinach

To gain a more detailed understanding of the formation and regulatory processes of the cavity, we performed single-cell transcriptome sequencing and analysis on the shoot tips of water spinach. We removed the leaves on shoot tips and harvested about 15 shoot tips for protoplast isolation (Supplementary Fig. 2b). To avoid the stress during protoplast isolation, we limited the time for protoplast isolation. We harvested the protoplasts at 0.5 h and 1 h after enzyme digestion separately, ensuring various cell types from shoot tips were isolated and captured (Supplementary Fig. 2c-d). The isolated protoplasts from both time points were combined and subjected to droplet-based scRNA-seq using the 10x Genomics scRNA-seq platform. In total, we obtained transcriptomic data from 19,181 single-cells. The median genes number per cell was 3,395 (Supplementary Table 3).

Unsupervised clustering identified 12 distinct cell clusters, and visualized using the uniform manifold approximation and projection (UMAP) algorithm and the t-distributed stochastic neighborhood embedding (t-SNE) tool (Fig. 2a, Supplementary Fig. 3). To assign cell identity to the clusters, we identified the marker genes of each cell cluster and examined the specificity of transcripts of known marker genes (Fig. 2c, Supplementary Fig. 4, and Supplementary Table 4-5). The 12 clusters covered all major cell types of the shoot tip, including epidermis, vasculature, photosynthetic cells, proliferating cells, meristem, and ground tissue (Fig. 2a-b). Epidermis group comprised of three clusters (Clusters 0, 6, 9), as indicated by epidermis-specific genes *protodermal factor 1* (*IaPDF1*), *glycosylphosphatidylinositol-anchored lipid protein transfer 1* (*IaLTPG1*), *eceriferum 3* (*IaCER3*), and *glycerol-3-phosphate sn-2-acyltransferase 1* (*IaGPAT1*) (Fig. 2c, Supplementary Fig. 4, Supplementary Table 5). Cluster 0 was identified as hypodermis since gene involved in lignin biosynthesis were highly expressed, such as, *caffeate o-methyltransferase 1* (*IaOMT1*), *cinnamoyl CoA reductase 1* (*IaCCR1*), and *peroxidase 52* (*IaPRX52*) (Fig. 2c, Supplementary Fig. 4, Supplementary Table 5).

**Fig 2.**
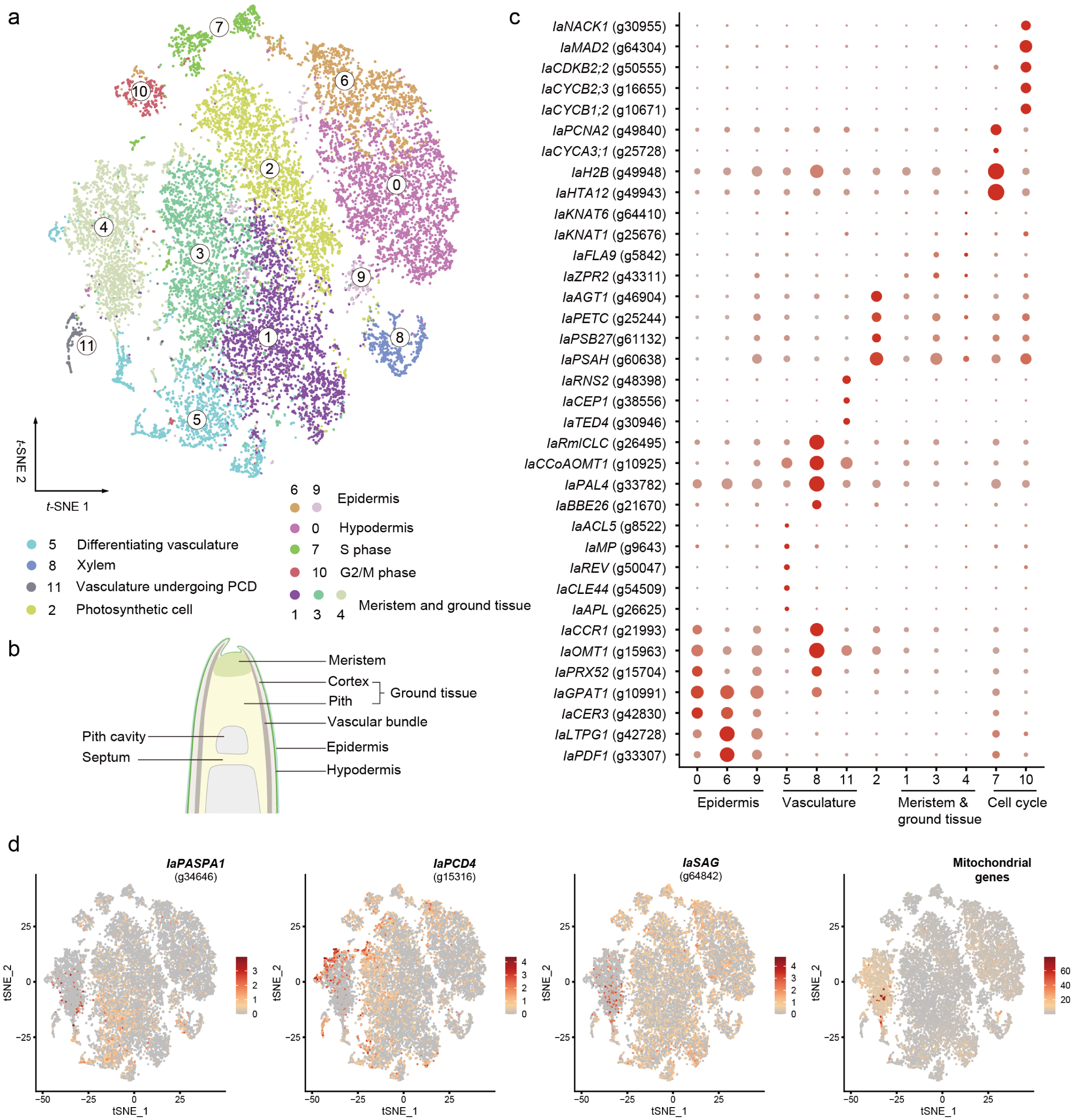
Cell atlas in the shoot tip of water spinach. a, Visualization of 12 cell clusters using t-SNE. dots, individual cells; n = 19,181 cells; color, cell clusters. b, Schematic of anatomy and cell types of shoot tip. c, Expression pattern of representative cluster-specific marker genes. dot diameter, proportion of cluster cells expressing a given gene. The full names of selected genes are given in Supplementary Table 5. d, t-SNE plot showing the selected PCD related genes and mitochondrial genes.

Vasculature group consisted of three clusters (Clusters 5, 8, 11), in which gene involved in xylem and phloem differentiation, and lignin biosynthesis were highly expressed (Supplementary Table 4). Cluster 5 was differentiating vasculature since gene involved in phloem formation (*altered phloem development*, *IaAPL*), cambial activity (*CLAVATA3/ESR-* related *44*, *IaCLE44*), and xylem specification (*REVOLUTA*, *IaREV*; *MONOPTEROS*, *IaMP*; *ACAULIS 5*, *IaACL5*) were markedly expressed (Fig. 2c, Supplementary Fig. 4, Supplementary Table 5). Cluster 8 was identified as more mature xylem since marker genes of xylem parenchyma were highly expressed, and some genes were involved in lignin biosynthesis, including *phenylalanine ammonia-lyase 4* (*IaPAL4*) and *caffeoyl coenzyme a o-methyltransferase 1* (*IaCCOAOMT1*) (Fig. 2c, Supplementary Fig. 4, Supplementary Table 5). Cluster 11 was likely vasculature undergoing PCD since both marker genes of xylem parenchyma (*IaCCOAOMT1* and *tracheary element differentiation-related 4*, *IaTED*) and PCD (cysteine endopeptidase 1, *IaCEP1*; *ribonuclease 2, IaRNS2*) were overrepresented (Fig. 2c, Supplementary Fig. 4, Supplementary Table 5). Cluster 2 comprises cells actively engaged in photosynthesis, in which genes involved in photosynthesis and photorespiration such as *photosystem I subunit H* (*IaPSAH*), *photosystem II subunit P* (*IaPSB27*), *photosynthetic electron transfer C* (*IaPETC*) and *alanine:glyoxylate aminotransferase 1* (*IaAGT1*) were predominantly expressed (Fig. 2c, Supplementary Fig. 4, Supplementary Table 5).

Clusters 7 and 10 were identified as proliferating cells because cell-cycle-related genes were overrepresented (Supplementary Table 4). Cluster 7 was assigned to S phase because it was enriched in chromatin organization or nucleosome assembly genes (*histone H2A 12*, *IaHTA12* and *histone H2B*, *IaH2B*), DNA biosynthesis (*proliferating cell nuclear antigen 2*, *IaPCNA2*), and marker gene of S phase (*cyclin A3;1*, *IaCYCA3;1*) (Fig. 2c, Supplementary Fig. 4, Supplementary Table 5). The expression of G2/M phase marker genes was restricted to cluster 10, such as *cyclin B 1;2* (*IaCYCB1;2*), *cyclin B2;3* (*IaCYCB2;3*), *cyclin-dependent kinase B2;2* (*IaCDKB2;2*), *mitotic arrest-deficient 2* (*IaMAD2*), and *NPK1-activating kinesin 1* (*IaNACK1*), suggesting that cluster 10 is undergoing mitotic cell division (Fig. 2c, Supplementary Fig. 4, Supplementary Table 5).

Clusters 1, 3, and 4 most likely compose the cell population corresponding to the area where the shift from a solid to a hollow pith occurred. Anatomically, pith cavity formed below the apical meristematic tissue. Clusters 1, 3, and 4 belong to meristem and pith, because the meristic marker genes (*knotted1-like homeobox gene 1*, *IaKNAT1* and *knotted1-like homeobox gene 6*, *IaKNAT6*) and pith marker genes (*Fasciclin-like arabinoogalactan 9*, *IaFLA9* and *LITTLE ZIPPER 2*, *IaZPR2*) were overrepresented (Fig. 2c, Supplementary Fig. 4, Supplementary Table 5). RNA *in situ* hybridization assays demonstrated that *aquaporin PIP2*, a gene specifically expressed in cell clusters 1, 3, and 4, was primarily localized in the meristem and its adjacent regions (Supplementary Fig. 5). These findings further confirmed that cell clusters 1, 3, and 4 are associated with the meristem-pith region. Furthermore, the expression of genes associated with PCD (*IaPASPA1*; *programmed cell death protein 4*, *IaPDCD4*; and *senescence-associated gene*, *IaSAG*), and mitochondrial gene levels progressively increases in clusters 1, 3, and 4 (Fig. 2d). The high content of mitochondrial genes is a characteristic of dying cells ^36^, and such genes are highly expressed in cluster 4 (Fig. 2d). It’s likely that clusters 1, 3, and 4 may have each experienced the process from the initiation of programmed cell death to execution leading to cell death.

### The developmental trajectory of pith cavity

To investigate the formation of pith cavity, we extracted meristem and pith cells (clusters 1, 3, and 4) under PCD by extracting cells with high expression level of three PCD associated genes (*IaPASPA1*, *IaPDCD4*, and *IaSAG*) (Fig. 2d). These cells were further unsupervisedly divided into five sub-cell clusters (Fig. 3a). We inferred the developmental trajectory using pseudotime analysis. In general, the trajectory started with sub-cluster 0 and end with sub-clusters 2 and 3, with two minor bifurcation points (Fig. 3b-c). To explore the developmental trajectory represented by pseudotime, we visualized PCD marker genes using UMAP plots (Fig. 3d). Despite different genes exhibit varying expression patterns across subclusters, overall, expression levels increased along with pseudotime (Fig. 3d). Additionally, expression of mitochondrial genes achieved its highest levels in subcluster 2, representing cells that were in the process of dying (Fig. 3d). Therefore, the pseudotime reflects the process of cavity development.

**Fig 3.**
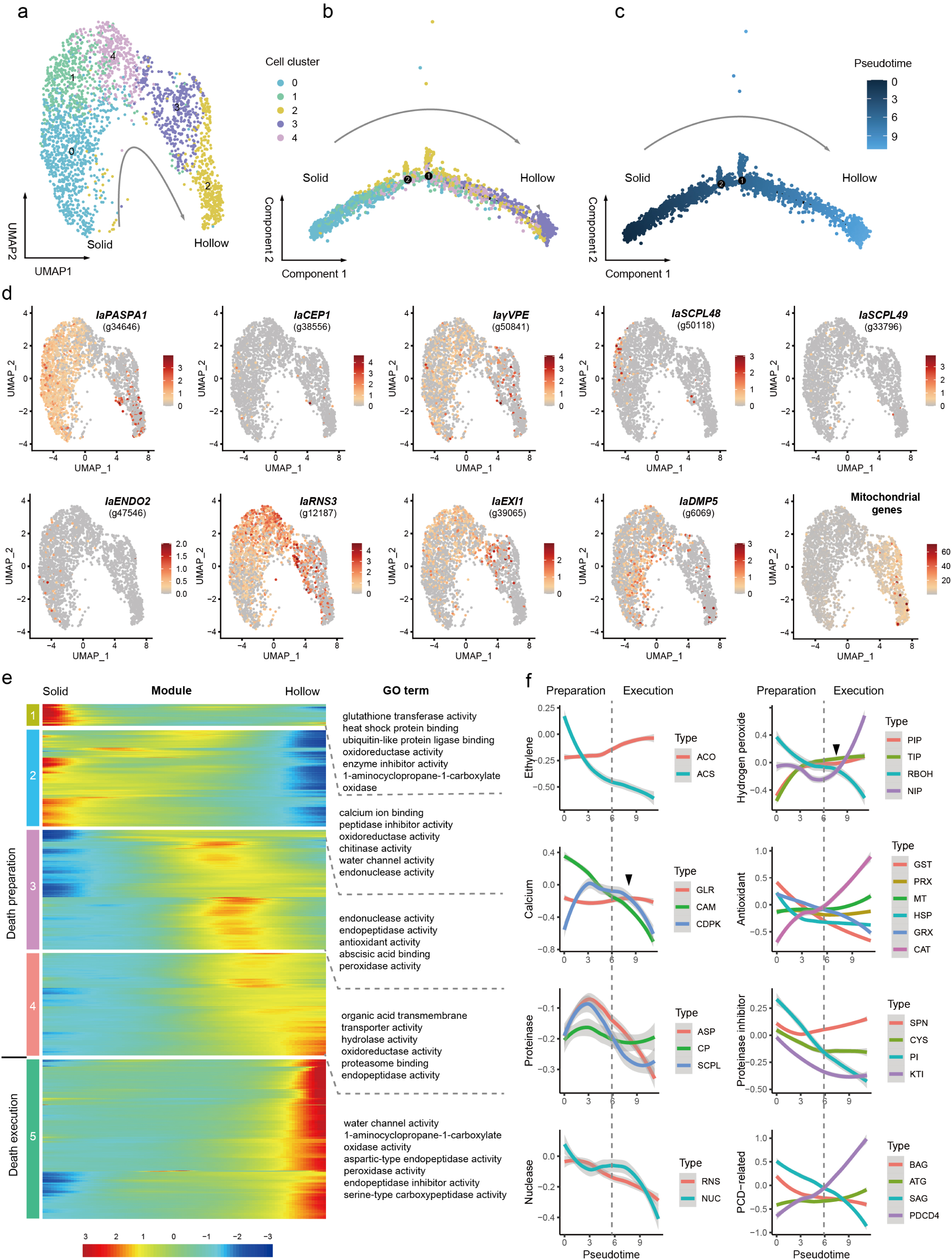
The developmental trajectory of pith cavity. a, Visualization of solid-hollow transition cell clusters using UMAP, colored by sub-cell clusters. n = 2,528 cells. b-c, differentiation trajectory from solid to hollow regions, colored by sub-cell clusters (b) or pseudotime (c). n = 2,528 cells. d, UMAP plot showing the expression pattern of PCD marker genes. e, Heat map showing the five gene modules of significant differentially expressed genes along pseudotime and their enriched GO terms. f, GSVA analysis showing the dynamic trends of significantly differentially expressed biological processes during cavity development along the pseudotime trajectory. ACS: ACC synthase, ACO: ACC oxidase, CAM: calmodulin, GLR: glutamate receptors, CDPK: calcium-dependent protein kinase, RBOH: respiratory burst oxidase homolog, AQP: aquaporin, PIP: plasma membrane intrinsic protein, TIP: tonoplast intrinsic protein, NIP: nodulin 26-like intrinsic protein, GST: glutathione S-transferase, HSP: heat shock protein, GRX: glutaredoxin, PRX: peroxidase, MT: metallothionein, CAT: catalase, ASP: aspartic proteinase, CP: cysteine proteinase, SCPL: serine carboxypeptidase-like protein, PI: proteinase inhibitor, CYS: cysteine proteinase inhibitor, KTI: kunitz trypsin inhibitor, SPN: serine protease inhibitor; RNS: ribonuclease, NUC: nuclease, BAG: B cell lymphoma 2 (Bcl-2) associated athanogene, SAG: senescence-associated gene, ATG: autophagy-related gene, PDCD4: programmed cell death protein 4.

To reveal the genes involved in spatial-temporal regulation of pith cavity formation, we identified 2,238 genes were significantly related to pseudo-time order (Supplementary Table 6). Genes that are significantly correlated with pseudotime can be clustered into five modules based on the patterns of their expression changes along the pseudotime trajectory (Fig. 3e). We performed GO enrichment analysis for each module. Generally, GO terms such as antioxidant activity, ethylene biosynthesis, protease activity, peptidase inhibition, endonuclease activity, and water channel activity exhibit changes in association with pseudotime (Fig. 3e). Based on pseudotime order, the modules 1-4 were most likely reflected death preparation, and module 5 was related to death execution.

Then, we selected the functional gene sets which significantly changed during the formation of pith cavity to calculate the gene set variation analysis (GSVA) and plotted the fit curve showing the relationship between GSVA values and pseudotime. Due to ethylene’s role in promoting PCD in plants, we observed the expression changes of ethylene biosynthesis genes. The expression of ACC synthase (ACS) was decreased as pseudotime, while ACC oxidase (ACO) expression increased along with pseudotime (Fig. 3f, Supplementary Fig. 6). Calcium signature and reactive oxygen species (ROS) are important signals to trigger PCD initiation and execution ^33,37^. The calmodulin (CAM), glutamate receptors (GLR) and calcium-dependent protein kinase (CDPK) are key actors of calcium sensing and signaling transduction ^38–40^. Whilst their expression patterns are different (Fig. 3f, Supplementary Fig. 6). The calcium sensor CAM decreased during the formation of pith cavity. The calcium mediator CDPK initially increased before subsequently decreasing. The expression of calcium channel GLR began with a slight decrease, followed by a minor increase, and then decreased again (Fig. 3f, Supplementary Fig. 6). A small second burst of CDPK and GLR could be recognized may be associated with death execution (Fig. 3f).

The calcium and CDPK can promote respiratory burst oxidase homolog (RBOH) to produce ROS ^41^. During the formation of pith cavity, expression level of RBOH generally decreased as pseudotime, but had a second burst during death execution (Fig. 3f, Supplementary Fig. 6). Aquaporins (AQPs) are water channels that facilitate transmembrane transport of H_2_O_2_ ^41,42^. Overall, the expression of two subfamilies of AQP, including plasma membrane intrinsic proteins (PIPs), tonoplast intrinsic proteins (TIPs) increased with pseudotime. While nodulin 26-like intrinsic proteins (NIPs) initially decreased, followed by an increase (Fig. 3f, Supplementary Fig. 6). Due to the ROS burst during the formation of pith cavity, a series of antioxidant pathways are activated to maintain the balance of the redox state. However, the spatiotemporal expression patterns of different antioxidant pathways varied (Fig. 3f, Supplementary Fig. 6). Glutathione S-transferases (GSTs), heat shock proteins (HSPs), glutaredoxins (GRXs) decreased along with pseudotime. The expression level of peroxidases (PRXs) changed slightly, initially decreasing before increasing again. While, metallothionein (MT) and catalase (CAT) significantly increased as pseudotime.

Proteases, protease inhibitors and nucleases directly regulate plant developmental PCD processes ^43^. Overall, aspartic proteinases (ASPs), cysteine proteinases (CPs) and a serine carboxypeptidase-like protein (SCPL) were accumulated initially, and decreased then. But during death execution, CPs and SCPLs increased slightly (Fig. 3f, Supplementary Fig. 6). Protease inhibitors bind to and inhibit proteases, thereby preventing precocious onset of PCD ^44,45^. Proteinase inhibitors (PIs), cysteine proteinase inhibitors (CYSs), kunitz trypsin inhibitors (KTIs) decreased as pseudotime, while serine protease inhibitors (SPNs) decreased initially and followed by an increase (Fig. 3f, Supplementary Fig. 6). Nucleases function directly in nuclear DNA degradation during PCD ^46,47^. The ribonucleases (RNSs) and nucleases (NUC) decreased during formation of pith cavity (Fig. 3f, Supplementary Fig. 6).

We also focused on other genes which have been reported involved in PCD processes. PDCD4 in plant, is a homologous gene of animal programmed cell death 4, has been implicated in ethylene signaling and PCD ^48,49^. The expression level of *IaPDCD4* increased during the formation of pith cavity (Fig. 3f, Supplementary Fig. 6). Plant PCD is also controlled by conserved protein family B cell lymphoma 2 (Bcl-2) associated athanogene (BAG) ^50^. Several BAG genes were significantly related to pseudotime order, and their expression levels decreased with pseudotime (Fig. 3f, Supplementary Fig. 6). Senescence-associated genes (SAGs) are involved in senescenece-asscociated programmed cell death ^51^, of which expression level decreased with pseudotime (Fig. 3f, Supplementary Fig. 6). Meanwhile, autophagy is required for timely cell death execution and corpse clearance ^52^. Autophagy-related genes (ATGs) increased during pith cavity formation (Fig. 3f, Supplementary Fig. 6).

### Identification of key TFs regulating cavity formation

By screening upregulated TFs during cavity development in the bulk and single-cell transcriptome, we found that NAC TFs were significantly enriched in both datasets (Supplementary Fig. 7). Consequently, we then screened NAC TFs for genes that potentially regulate PCD or cell death based on the expression patterns. By analyzing the expression trends along the pseudotime trajectory, we focused on genes whose expression levels increased along with pseudotime, particularly in the end of pseudotime trajectory (death execution). Combined with previous functional reports, we identified three NAC TFs (*IaNAC074*, *IaNAC087* and *IaNAC029*) that were significantly upregulated in the regions undergoing cell death and play potential roles in cell death (Fig. 4a). Subcellular localization assays revealed that these NAC TFs were localized in the nucleus (Fig. 4b).

**Fig 4.**
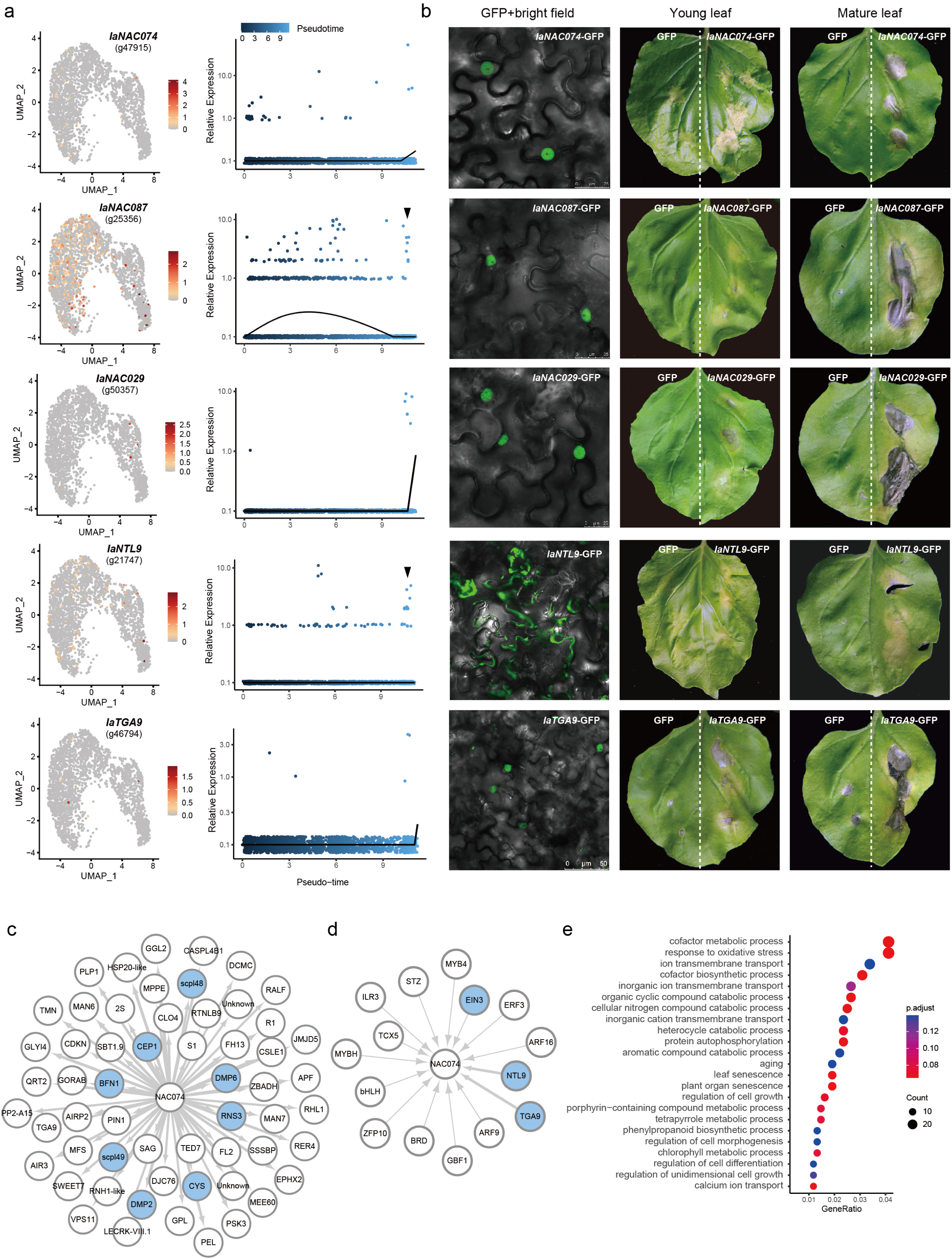
Regulatory TFs involved in cavity formation. a, Expression pattern of regulatory TFs in UMAP plots and along pseudotime trajectory. b, Subcellular localization and phenotypic observations following TFs transiently expressed in *Nicotiana benthamiana* leaves. Agrobacteria containing different constructs were inoculated in leaves, and GFP signal was imaged at 2 days after inoculation using a confocal microscope. The leaves expressing *pro35S::IaNAC074-GFP*, *pro35S::IaNAC087-GFP*, *pro35S::IaNAC029-GFP*, *pro35S::IaTGA9-GFP*, and *pro35S::IaNTL9-GFP* were taken photos using a digital camera at 12 days after inoculation. The *pro35S::GFP* vector served as a negative control. c, Gene-regulatory network of *IaNAC074* as regulator, inferred by gene that are dynamically expressed across pith cavity development based on scRNA-seq data. Blue node highlights PCD-related genes (interaction weight>0.025). Edge width indicates the interaction weight. d, Gene-regulatory network of *IaNAC074* as target (interaction weight>0.01). e, GO enrichment analysis of *IaNAC087*-regulated genes predicted using GEINIE.

*IaNAC074* exhibited the highest expression levels at the end of the pseudotime trajectory (Fig. 4a). To investigate its function, we transiently expressed *IaNAC074* in *Nicotiana benthamiana* leaves. Within 1-2 days post-injection, cell death was observed in the injected sites on tobacco leaves. In young leaves, the injected regions turned yellow and wilted, while the injection area in mature leaves became transparent (Fig. 4b, Supplementary Fig. 8). Notably, the cell death phenotype was confined only to the injected regions. Furthermore, the gene regulatory networks of cavity-developing cells suggested that *IaNAC074* regulates multiple downstream genes involved in death execution, such as *IaCEP1*, *IaRNS3*, *IaBFN1*, *IaDMP2*, *IaDMP6*, and *IaSCPL49* (Fig. 4c). GO enrichment analysis of these genes regulated by *IaNAC074* also revealed a significant association with PCD (Supplementary Table 7). Additionally, transient expression of *IaNAC074* in tobacco leaves resulted in increased expression of death execution genes, such as *NbPASPA1*, *NbCEP1*, *NbBFN1*, *NbDMP2*, and *NbSCPL48* (Fig. 5e). Previous studies reported that *NAC074* controls the expression of PCD–associated genes, such as *CEP1*, *RNS3*, *BFN1* and *EXI1*, and promotes cell death in the stigma of Arabidopsis and the stems of sorghum ^11,28^. Therefore, all current evidences suggest that *IaNAC074* is a key regulator of PCD during cavity development in water spinach.

**Fig 5.**
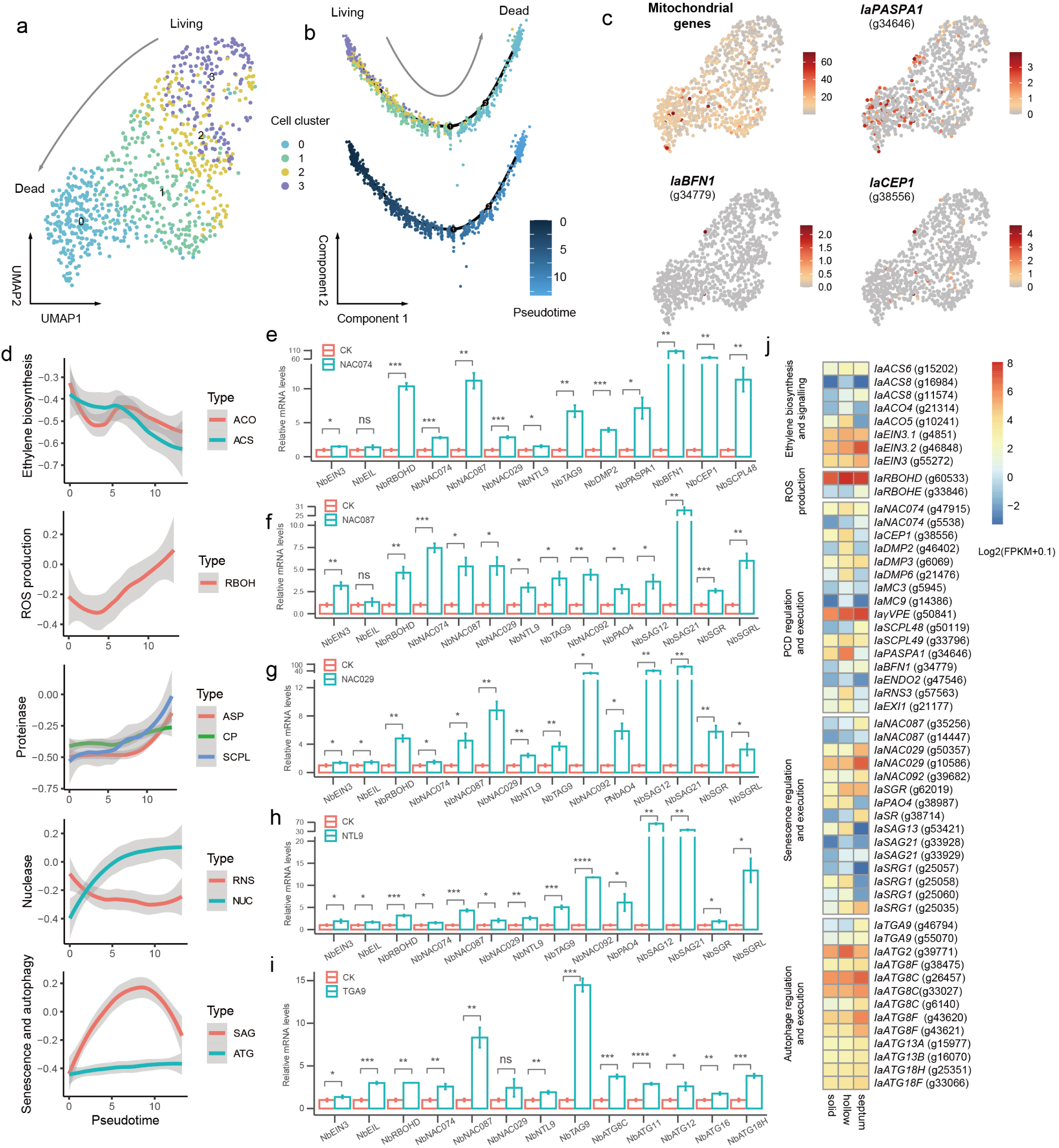
PCD, senescence, and autophagy collaboratively contribute to death execution. a, Visualization of dying cell clusters using UMAP, colored by sub-cell clusters. n = 861 cells. b, differentiation trajectory from cell death, colored by sub-cell clusters (top) or pseudotime (bottom). n = 861 cells. c, UMAP plot showing the expression pattern of PCD marker genes. f, GSVA analysis showing the dynamic trends of significantly differentially expressed pathway during cell death along the pseudotime trajectory. ACS: ACC synthase, ACO: ACC oxidase, RBOH: respiratory burst oxidase homolog, ASP: aspartic proteinase, CP: cysteine proteinase, SCPL: serine carboxypeptidase-like protein; RNS: ribonuclease, NUC: nuclease, SAG: senescence-associated gene, ATG: autophagy-related gene. e-i, RT-qPCR validation of downstream genes regulated by *IaNAC074*, *IaNAC087*, *IaNAC029*, *IaNTL9*, and *IaTGA9* by transient expression in tobacco leaves, with empty vector as control. Statistical significance was calculated using t test. ns: not statistically significant, *: p <= 0.05, **: p <= 0.01, ***: p <= 0.001, ****: p <= 0.0001. j, Heat map of gene expression based on bulk RNA transcriptome in solid, hollow regions and septum of shoot tips, sampled regions illustrated in Supplementary Fig. 1a.

*IaNAC087* was expressed throughout the hollowing process, with increased expression observed at the end of the pseudotime trajectory (marked by the black arrow in Fig. 4a). After the injection of *IaNAC087* into tobacco leaves, leaf yellowing occurred within 5-6 days. In mature leaves, cell death was observed in the injected regions after 9-10 days, with yellowing extending beyond the injected regions (Fig. 4b, Supplementary Fig. 8). *ANAC087* plays a significant role during PCD in Arabidopsis roots, although it does not directly regulate PCD-execution genes *in vivo* ^30^. In water spinach, GO enrichment analysis of *IaNAC087*-regulated genes indicated a significant association with leaf senescence in cavity-developing cells (Fig. 4e, Supplementary Table 8). Quantitative transcriptional analysis confirmed that transient expression of *IaNAC087* in tobacco leaves enhanced the expression of senescence-related genes, including *NbNAC092*, *NbPAO4*, *NbSAG12*, *NbSAG21*, *NbSGR*, and *NbSGR-like* (Fig. 5f). In addition, the transient expression of *IaNAC087* in tobacco also resulted in senescence phenotypes, indicating that *IaNAC087* is likely induce cell death by triggering senescence, at least in water spinach.

Senescence contributes to cell death in plants. In Arabidopsis, *ANAC092* (also named as *ORESARA1*) is a well-known regulator of senescence and is involved in cell death of stigma. During the cavity development in shoot tips of water spinach, *IaNAC092* was upregulated at the late stages of cavity development (Supplementary Fig. 9). In the meantime, another prominent senescence-associated TF, *NAC029*, was found to be exclusively expressed at the late stages of cavity development (Fig. 4a). After injection of *IaNAC029* into tobacco leaves, yellowing occurred within 4-5 days, followed by gradual cell death in the injected regions both in young and mature leaves (Fig. 4b, Supplementary Fig. 8). Transient expression of *IaNAC029* in tobacco leaves resulted in increased expression of senescence-associated genes (including *NbNAC092*, *NbPAO4*, *NbSAG12*, *NbSAG21*, *NbSGR*, and *NbSGR-like*) (Fig. 5g). In cavity-developing cells, GO enrichment analysis revealed that leaf senescence was significantly enriched in *IaNAC029*-regulated genes (Supplementary Table 9). These indicate that *IaNAC029* likely promotes cell death in shoot tips of water spinach by inducing senescence.

In order to identify more regulators of cell death in pith, we utilized *IaNAC074* as a bait gene to search the gene regulatory networks to identify possible regulators of *IaNAC074* or co-expressed TF. We found *IaNTL9* (NAC with Transmembrane Motif 1-like TF 9) and *IaTGA9* (TGACG motif-binding protein 9) are two most closely related genes to *IaNAC074* (interaction weight of links as 0.034 and 0.030 respectively) (Fig. 4d). *IaNTL9* localized on the cell membrane (Fig. 4b), and it was expressed throughout the hollowing process but showed elevated expression in dying cells (marked by the black arrow in Fig. 4a). Injection of *IaNTL9* into tobacco leaves caused leaf yellowing within 5-6 days but did not lead to cell death in both young and mature leaves (Fig. 4b, Supplementary Fig. 8). Transient expression of *IaNTL9* in tobacco leaves resulted in increased expression of senescence-associated genes, such as *NbNAC092*, *NbPAO4*, *NbSAG12*, *NbSAG21*, *NbSGR*, and *NbSGR-like* (Fig. 5h). NTL9 is involved in various physiological processes, including abiotic and immune signaling responses, and regulating developmental leaf senescence, as reported in previous studies ^53^. In cavity-developing cells of water spinach, leaf senescence was also significantly enriched in *IaNTL9*-regulated genes (Supplementary Table 10). Therefore, *IaNTL9* contributes to cell death regulation by inducing senescence.

*TGA9* is an important autophagy-related TF that executes cell autophagy through *ATG8* (autophagy-related 8) gene ^54^. Subcellular localization assay revealed that *IaTGA9* was localized in the nucleus (Fig. 4b). *IaTGA9* expression increased at the late stages of cavity development (Fig. 4a). Injection of *IaTGA9* into tobacco leaves resulted in leaf yellowing within 5-6 days and eventual death of both young and mature leaves (Fig. 4b, Supplementary Fig. 8). Quantitative transcriptional analysis confirmed that transient expression of *IaTGA9* in tobacco leaves enhanced the expression of *ATG* genes, including *NbATG8C*, *NbATG11*, *NbATG12*, *NbATG16*, and *NbATG18H* (Fig. 5i). As revealed by single-cell transcriptome data of water spinach, autophagy was significantly enriched in *IaTGA9*-regulated genes (Supplementary Table 11). These findings indicate that *IaTGA9* promotes cavity formation by regulating autophagy.

### PCD, senescence, and autophagy collaboratively contribute to cell death

To more precisely investigate the gene regulation during the process of cell death, we isolated death execution related cells (subclusters 2 and 3 of solid-hollow transition cells in Fig. 3a) for further cellular clustering and pseudotime analysis. These cells were unsupervisedly divided into four sub-cell clusters (Fig. 5a). The trajectory started with sub-cluster 3 and end with sub-clusters 0 (Fig. 5b). PCD marker genes, including *IbPASPA1*, *IbBFN1* and *IaCEP1*, and mitochondrial genes were highly expressed in sub-cell clusters 0 and 1 (Fig. 5c). Therefore, the pseudotime represents the process of cell death.

In total, 1,559 genes were identified that significantly related to pseudo-time order (Supplementary Table 12). Then, we calculate GSVA values for the selected genes and plotted the fit curves showing the relationship between GSVA values and pseudotime (Fig. 5d). The two ethylene biosynthesis genes showed a decline, both with a slight increase in the middle. The overall expression trend of RBOHD was increased along with pseudotime. The expression of proteases, including ASPs, CPs and SCPL, were significantly increased during cell death. As for nuclease, NUC increased during cell death, while RNS decreased. The expression of SAG increased initially but decreased with pseudotime. ATG constantly increased during cell death. Therefore, analysis of dying cell clusters indicates that genes involved in PCD, senescence, and autophagy collaboratively participate in cell death. Additionally, bulk transcriptome data also reveal that during cavity development and cell death, there is an upregulation of genes associated with ethylene synthesis and signaling, ROS production, PCD, senescence, and autophagy (Fig. 5j).

### Ethylene triggered pith cavity in shoot tips

Since genes involved in ethylene biosynthesis (*IaACS* and *IaACO*) exhibited significant changes during cavity formation and cell death (Fig. 3f, Fig. 5d), we hypothesized that ethylene may be one of the plant hormones inducing cavity formation in shoot tips of water spinach. To test this, we cultured shoot tips on MS media supplemented with the ethylene synthesis inhibitor aminoethoxyvinylglycine (AVG) and the ethylene precursor 1-aminocyclopropanecarboxylic acid (ACC). Shoot tips grown on AVG-supplemented media showed growth arrest phenotype, with no significant shoot elongation and barely development of new internodes (Fig. 6a-c). In contrast, shoot tips grown on ACC-supplemented media showed a significant increase in plant height and new internode number compared to the control (Fig. 6a-c). These results suggested that ethylene promotes both node formation (related to cavity formation) and internode elongation, while the absence of ethylene inhibits these processes.

**Fig 6.**
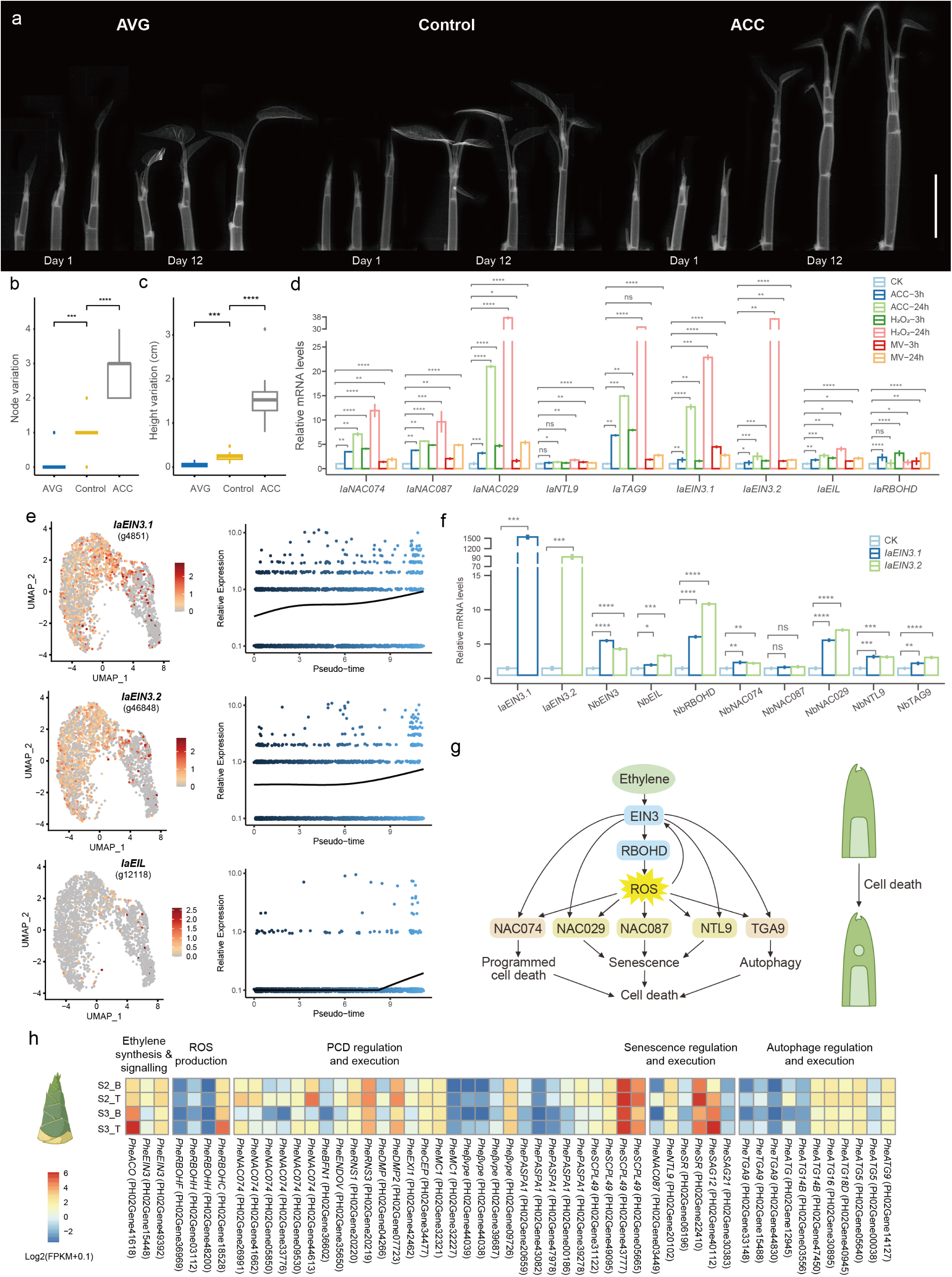
Ethylene triggered cavity formation in shoot tips. a, The phenotypic differences among shoot of water spinach on MS medium supplemented with AVG and ACC, and untreated MS medium as the control. Scale bar, 1 cm. b-c, The variation of node number (b) and height (c) of shoots under the treatments of AVG and ACC compared with control. The results are derived from three biological repetitions. Statistical significance was calculated using wilcoxon test. ***: p <= 0.001, ****: p <= 0.0001. d, The relative expression of genes in water spinach leaves at 3 and 24 h after treatments of ACC, H_2_O_2_, and MV, with water treatment after 24h used as the control. Statistical significance was calculated using t test. ns: not statistically significant, *: p <= 0.05, **: p <= 0.01, ***: p <= 0.001, ****: p <= 0.0001. e, Expression pattern of *IaEIN3* and *IaEIL* in UMAP plots and along pseudotime trajectory. f, The relative expression level of identified death-regulating genes in tobacco leaves infiltrated with Agrobacterium harboring *pro35S::IaEIN3.1* and *pro35S:: IaEIN3.2*, respectively. Statistical significance was calculated using t test. g, Molecular regulatory mechanisms of pith cavity formation in water spinach. h, Heat map of gene expression in the apical and basal internodes of moso bamboo (*Phyllostachys edulis*). S2: rapid growth stage, spring shoots are approximately 50 cm high aboveground; S3: slow growth stage, spring shoots are approximately 150 cm high aboveground; T: apical internode; B: basal internode.

To further verify the role of ethylene in promoting cavity formation, we sprayed ACC solution on the leaves of water spinach and examined the expression changes of key regulatory genes involved in cavity development. The results showed that ethylene treatment led to increased expression of RBOHD and upregulated death-regulating TFs (*IaNAC074*, *IaNAC087*, *IaNAC029*, *IaNTL9*, *IaTGA9*) (Fig. 6d). This suggests that ethylene may induce cavity formation by regulating ROS production and directly modulating death-regulating TFs. To confirm the role of ROS in promoting cavity formation, we applied hydrogen peroxide and methyl viologen to mimic exogenous and endogenous ROS accumulation, respectively. The results demonstrated that both exogenous and endogenous ROS elevated the expression of five death-regulating TFs, and EIN3 (Ethylene Insensitive 3) and EIN3-Like (EIL), which are the central TFs in ethylene signaling (Fig. 6d). These findings indicated that ethylene and ROS interact synergistically to promote cavity formation by inducing the expression of death-regulating TFs.

Since EIN3 is the central TF in ethylene signaling, we hypothesize that ethylene may regulate cavity development through EIN3. In water spinach, the expression of *IaEIN3.1* (g4851), *IaEIN3.2* (g46848), and *IaEIL* (g12118) indeed increased during cavity development. Notably, both *EIN3* genes have shown extremely high expression levels (Fig. 6e). Gene regulatory network analysis based on single-cell transcriptomics also indicates that *IaEIN3* is a regulatory gene of *IaNAC074* (Fig. 4d). These suggested that ethylene may regulate cavity formation through EIN3. To test this, we employed an *Agrobacterium*-mediated transient expression system in *N. benthamiana*. Compared to the empty vector control, transient expression of *IaEIN3.1* or *IaEIN3.2* led to a significant increase in *IaEIN3.1* and *IaEIN3.2* expression respectively. Concurrently, genes and TFs involved in cell death regulation also showed varying levels of upregulation, with *NbRBOHD* and *NbNAC029* expression increasing by 5-11 times and *NbNAC074*, *NbTGA9* and *NbNTL9* by 2-3 times, while *NbNAC087* showed minimal upregulation (Fig. 6f). These results suggested that ethylene regulates *RBOHD* and most of death-regulating TFs (*NAC074*, *NAC029*, *NTL9* and *IaTGA9*) through EIN3, but the regulation of *NAC087* may be independent from EIN3.

### Evolutionary conservation of stem cavity formation in plants

The bamboo culm, is also segmented with distinct nodes and hollow internodes. During the development of moso bamboo (*Phyllostachys edulis*) shoots, the apical internodes are undergoing cavity formation, while the basal internodes have already completed this process. By comparing gene expression differences between the apical and basal internodes, we found that genes and TFs involved in ethylene signaling, the regulation and execution of PCD, senescence, and autophagy were upregulated in the apical internodes (Fig. 6h). This suggested that these pathways also play a role in the formation of the pith cavity in bamboo. Previous studies on cavity development in bamboo have also shown that ethylene signaling pathways, RBOH and PCD-related genes are upregulated during the process of cavity formation ^20^. The hollow stem formation in the dicotyledonous water spinach and the monocotyledonous moso bamboo represents a parallel evolutionary event, whilst the underlying mechanisms are similar, reflecting the evolutionary conservation of precise cell death regulation in plants.

## Discussion

Hollow stems are often associated with specific environmental adaptations, such as survival in aquatic or waterlogged conditions, as seen in water spinach and reed (*Phragmites australis*). The identification of ethylene and ROS as key drivers in this process suggests that plants may have developed a sophisticated response to environmental pressures, using these signaling molecules to modify their internal structure for survival. The pith cavity in water spinach is initiated by ethylene and ROS, which then activate pathways involved in PCD, senescence, and autophagy, leading to pith cell death (Fig. 6g). As the shoot tip elongates, the cavity gradually expands. During the initial stage, ethylene and ROS signaling mutually enhance each other, jointly regulating TFs that trigger PCD, senescence, and autophagy, such as, *IaNAC074*, *IaNAC087*, *IaNAC029*, *IaNTL9*, and *IaTGA9* (Fig. 6g). Calcium signaling may also play a critical role throughout the cavity formation process. As the development of the cavity, we observed dynamic changes in the expression of calcium sensing and signaling genes, such as calmodulins, calcium channel and calcium-dependent protein kinase. These findings are consistent with previous reports suggesting that calcium signaling is crucial in regulating various developmental PCD in plants ^33^. In the development of the pith cavity, cell death is precisely controled via PCD, while genes associated with senescence and autophagy facilitate the degradation and recycling of cellular materials.

The pith cavity formation in dicot water spinach and monocot bamboo represents a case of parallel evolution, where both species utilize similar molecular mechanisms for cavity development. Despite the evolutionary distance between dicots and monocots, there is a convergent use of pathways governing PCD for the development of structural features such as hollow stem. By further comparing the mechanisms of pithiness across different plant species, we uncovered a strong evolutionary conservation in the precise regulation of cell death. Precise regulation of cell death during plant development is crucial, a process also known as developmental programmed cell death (dPCD). Plant hormones often act as initiators, triggering the cell death process through TFs ^33^. Our study found that during the during the execution phase of cell death, the expression of ethylene synthesis genes gradually decreases, confirming the role of ethylene in initiating the cell death process. Subsequently, ROS act as secondary messengers, initiating and regulating PCD ^33^. Our study found that ethylene and ROS can mutually enhance each other, thereby co-regulating the process of cell death. Ethylene and ROS also act as key regulators in anther development, the formation of aerenchyma in the stems of aquatic plants and in the roots during flooding ^26,27,33^. Programmed cell death in different cell types is precisely regulated by distinct TFs. *NAC074* is likely a regulatory factor involved in the death of pith cells. Because *NAC074* is not only essential for the cavity formation in water spinach, but also plays a role in pith death in sorghum stems and in the senescence of the Arabidopsis stigma ^11,28^. While the role of senescence in developmental PCD has often been overlooked. Senescence is considered a form of PCD, referred to as senescence-associated PCD^51^. The senescence-associated TFs *NAC092* and *NAC074* jointly regulate the process of stigma senescence in Arabidopsis. In this study, we found that the formation of stem cavities in both water spinach and bamboo involves senescence in conjunction with PCD. These results further supporting that senescence is tightly integrated with PCD regulation. Autophagy is reported as a downstream mechanism for clearance of terminally cell corpses during developmental PCD ^55,56^. Our study also indicates that autophagy plays a crucial role in the later stages of cell death, as expression levels of ATGs increase as the cell death process.

The ability to control hollow stem formation could have practical agricultural applications. Hollow stems provide certain advantages in plant architecture, such as reduced weight and better mechanical support in specific conditions, which can be useful in crop plants growing in flooded areas or wetlands. The knowledge gained from this study could be used to manipulate ethylene and ROS signaling pathways to engineer crops that are better adapted to waterlogged or poorly oxygenated soils. This could lead to the development of more resilient crops with optimized growth and productivity in challenging environments, particularly as climate change increases the frequency of extreme weather events like floods. In addition, the identification of specific TFs (*IaNAC074*, *IaNAC087*, *IaNAC029*, *IaNTL9*, and *IaTGA9*) involved in PCD, senescence, and autophagy provides molecular targets for genetic manipulation. By fine-tuning the expression of these factors, we could potentially control the timing and extent of cell death, allowing for the development of new crop varieties with hollow stems or other beneficial structural features. For example, this approach could be used to create plants with improved nutrient transport, aeration, or drought resistance, expanding the range of environments where crops can be cultivated successfully.

In addition, the suggestion that the same regulatory mechanisms may govern hollow stem formation in bamboo points to a broader conservation of death-regulatory pathways in plants. This finding indicates that other species with hollow stems, such as rice, certain grasses, and aquatic plants, may share similar molecular processes. Understanding this conserved pathway opens new avenues for cross-species research, where discoveries in one plant can inform genetic or physiological interventions in another. This discovery will accelerate advancements in plant science and agriculture by enabling the application of knowledge across a wide range of species.

## Method

### Plant materials and growth conditions

The water spinach used in this study was a leafy vegetable variety collected from Songjiang District, Shanghai. The seed was sterilized and transferred to MS medium (MS 4.4 g/L, sucrose 30 g/L, Gelrite 3 g/L, pH 5.8). The plant then propagated via tissue culture techniques and maintained under 16 hours light and 8 hours dark at 25°C.

### Morphological assays

#### X-ray scanning

The shoots of water spinach were placed in square culture dishes for cultivation. The development of the shoot tips was monitored daily using an X-ray machine (Faxitron UltraFocus60 X).

#### Histochemical staining

The shoot tips were longitudinally bisected. 0.4% trypan blue staining solution (Solarbio, Cat No./ID: C0040) and 0.5 mg/ml DAB solution (Sigma-Aldrich, Cat No./ID: D12384) was then applied to vertical-sections of shoot tip, ensuring that the stain covered the cut surfaces and filled the cavities. The tissues were stained for 10 minutes and then observed under a stereomicroscope (Nikon SMZ 1500).

#### Paraffin section

The shoot tips were harvested and fixed with FAA (50% ethanol, 5% glacial acetic acid, 3.7% formaldehyde and 41.3% water). Tissues were dehydrated with a series of ethanol treatments (70%, 90%, 95%, 100% and 100%), cleared with xylene, and embedded with paraplast (Solarbio, Cat No./ID: YA0011). Serial transverse sections (10 μm thick) obtained from paraffin embedded tissues using a microtome (Leica, RM2265). The resulting sections were then dewaxed for staining, TUNEL assay and RNA *in situ* hybridization. Sections were stained with toluidine blue solution (Solarbio, Cat No./ID: G4807) and examined with a microscope (Zeiss Axio Scope. A1). TUNEL assay was carried out according to the instructions of Colorimetric TUNEL Apoptosis Assay Kit (Beyotime, Cat No./ID: C1091). Subsequently, sections were observed under a microscope (Zeiss Axio Scope. A1).

### RNA *in situ* hybridization

DIG-EASY-BYB buffer solution was preheated at 37℃ and soaked the glass slides for 30 minutes, after that the glass slides were taken out and soaked with DIG-EASY-BYB buffer solution with probe for 18 hours at 42℃. The probe preparation was prepared according to ROCHE instructions, the probe concentration was set as 10-50 pg/ml. The slides were rinsed at room temperature with 2 X SSC twice, followed by rinsed at 65 ℃ with 0.5x SSC twice, 30 minutes per sessions. The slides were then rinsed in a washing buffer for 5 minutes and incubated in blocking solution for 2 hours, after that the slides were incubated in antibody solution at 37 ℃ for 2 hours. The glass slides were washed in a washing buffer twice with 15 minutes per sessions, and balanced in detection buffer for 5 minutes, the slides were put into the substrate color solution to avoid light inoculation.

### Genome assembly

The leaves of water spinach were collected for Nanopore sequencing, yielded 24.31 Gb sequencing data. The reads were assembled by NextDenovo (Version 2.3.0). The contigs were correct with Illumina sequencing reads by NextPolish (Version 1.2.3). 3D-DNA pipeline was used to anchor the corrected contigs to the pseudo chromosomes and generate a Hi-C heatmap. For errors in the Hi-C heatmap, the Juicebox assembly Tools module was employed to identify and correct manually (Fig. 1q). BUSCO v5.4.3 was used in each step to assess the quality of the assembled and polished genome. Gene structure annotation was predicted using AUGUSTUS (Version 3.3.3) ^57^. To annotate the predicted genes, we used BLASTx program (http://www.ncbi.nlm.nih.gov/BLAST/) with an E-value threshold of 1e-5 to NCBI non-redundant protein (Nr) database (http://www.ncbi.nlm.nih.gov), the Swiss-Prot protein database (http://www.expasy.ch/sprot), the Kyoto Encyclopedia of Genes and Genomes (KEGG) database (http://www.genome.jp/kegg), and the COG/KOG database (http://www.ncbi.nlm.nih.gov/COG). Gene annotations were subsequently obtained according to the best alignments.

### RNA-seq analysis

Total RNA was isolated from mixed shoot tips of water spinach and sweetpotato. A total amount of 2 μg RNA per sample was used as input material for the RNA sample preparations. Sequencing libraries were generated using the NEBNext® Ultra™ RNA Library Prep Kit for Illumina® (#E7530L, NEB, USA). The RNA concentration of sequencing library libraries were measured using Qubit 3.0 and insert size was assessed using the Agilent Bioanalyzer 2100 system (Agilent Technologies, CA, USA). The libraries were sequenced on an Illumina platform (Novaseq 6000) and 150 bp paired-end reads were generated. The transcriptome data of moso bamboo was downloaded from NCBI (PRJNA689987); further details can be found in the article ^58^.

The raw data were first processed with FastQC (http://www.bioinformatics.babraham.ac.uk/projects/fastqc/) to remove adapters and low-quality sequences. The RNA-seq reads were mapping against the assembled reference genome of water spinach using HISAT2 version 2.1.0^59^. StringTie version 2.1.4 ^60^ was employed to calculate FPKM (fragments per kilobase of transcript per million fragments mapped) of genes for each sample. The expression matrix was extracted by R package Ballgown ^61^. The expression heat map was visualized using R package pheatmap ^62^. Differential gene expression analysis was performed using the R software package DESeq2 ^63^. GO enrichment of selceted genes was performed by R package clusterProfiler ^64^.

### scRNA-Seq library construction and sequencing

To perform the scRNA-seq, water spinach shoot tips (approximately 1 cm in length) were collected and their leaves were removed. About 15 shoot tips were digested in enzyme solution: 1% (w/v) Cellulase Onuzuka R-10, 0.3% (w/v) macerozyme R-10, 0.2% (w/v) pectinase Y-23, 0.4 M Mannitol, 20 mM MES [2-(N-Morpholino) ethanesulfonic acid hydrate], 20 mM KCl, 10 mM CaCl_2_ and 0.1% bovine serum albumin (BSA), Ph 5.5∼5.8. After gentle shanking for 1 hour at 28 ℃ in the dark, the protoplasts were filtered out with 100 µm filter and concentrated by centrifugation (200 g for 5 min). The up part liquid was discarded and collected protoplasts were washed with 8% mannitol for three times and followed by filtering twice with 40 µm filter. Protoplast viability was determined by trypan blue staining and fluorescein diacetate staining for the purpose of quality inspection, the viable cells rate was about 90%. The protoplast concentration was adjusted to around 1000 cells/µl using 8% mannitol.

Around 20,000 cells were initially mixed with 10X Genomics single-cell reaction reagents for the scRNA-seq assay. Cellular suspensions were loaded on a 10X Genomics GemCode Single-cell instrument that generates single-cell Gel Bead-In-EMlusion (GEMs). Libraries were generated and sequenced from the cDNAs with Chromium Next GEM Single Cell 3’ Reagent Kits v3.1. Upon dissolution of the Gel Bead in a GEM, primers containing (i) an Illumina® R1 sequence (read 1 sequencing primer), (ii) a 16 nt 10X Barcode, (iii) a 10 nt Unique Molecular Identifier (UMI), and (iv) a poly-dT primer sequence were released and mixed with cell lysate and Master Mix. The Single Cell 3’ Protocol produced Illumina-ready sequencing libraries. A Single Cell 3’ Library comprised standard Illumina paired-end constructs which begin and end with P5 and P7. The Single Cell 3’ 16 bp 10X Barcode and 10 bp UMI were encoded in Read 1, while Read 2 was used to sequence the cDNA fragment. Sample index sequences were incorporated as the i7 index read. The cDNA libraries were sequenced on the Illumina sequencing platform by Majorbio Bio-pharm Technology Co., Ltd (Shanghai, China).

### scRNA-seq data analysis

The raw files were analyzed by Cell Ranger 3.0.1(10x Genomics) under default settings using our newly assembled genome and published chloroplast (NC_056300.1) and mitochondrial genome (MZ240730.1) as reference. The gene-cell matrices (named ‘filtered_gene_bc_matrices’ by 10x Genomics) were subsequently processed and filtered using Seurat (version 4.0.3 with R version 4.0.2; Stuart et al., 2019). The low-quality cells and genes were filtered as follows: (1) only the cells in which numbers of expressed genes was 200 to 7,000 were considered; (2) the cells with unique molecular identifiers (UMIs) above 30,000 were filtered out; (3) the genes that were expressed in fewer than 3 cells were removed. After quality control, normalization, and scaling, highly variable genes were identified, and principal component analysis (PCA) was performed for dimensionality reduction. The first ten significant principal components were used for clustering the cells using a graph-based clustering approach. Clusters were visualized using UMAP (Uniform Manifold Approximation and Projection) and t-SNE (t-distributed Stochastic Neighbor Embedding) plots. Marker genes for each cell cluster were identified using the FindMarkers function. The mitochondrial gene percentage was calculated using all mitochondrial genes. Cell populations involved in cavity development were extracted based on the expression levels of death-related genes (*IaPASPA1* > 0 or *IaSAG* > 2 or *IaPCD4* > 1).

To understand the developmental trajectories of the cells, pseudotime analysis was performed using the Monocle2 package (version 2.16.0). The preprocessed data from Seurat were imported into Monocle2, and cells were ordered in pseudotime to infer developmental trajectories. Differential gene expression analysis along the pseudotime trajectory was conducted to identify genes associated with specific states or transitions.

To integrate the expression trends of multiple copy genes, Gene Set Variation Analysis (GSVA) was performed using R package GSVA_1.36.3. GSVA was used to assess pathway enrichment and expression changes across the cells. For visualizing the pseudotime expression trends, the smoothed expression trajectories were plotted using R package ggplot2. Additionally, the GSVA results and pseudotime trends were projected onto a UMAP plot for better visualization of the spatial and temporal expression patterns using FeaturePlot function in Seurat. The gene regulatory networks was inferred using GENIE3_1.10.0 ^65^.

### Transient expression assay and subcellular location

The coding sequence (CDS) was ligated into the vector pCAMBIA1302 and GFP was fused to the C-terminals of *IaNAC074*, *IaNAC087*, *IaNAC029*, *IaTGA9*, and *IaNTL9*, respectively. The CDS of *IaEIN3.1* and *IaEIN3.2* were cloned into the vector pGreenII 62-SK. The constructs and empty vectors were introduced into *Agrobacterium tumefaciens* strain GV3101 for Agrobacterium-mediated transient transformation. Overnight cultures of *A. tumefaciens* cells were centrifuged and suspended in infiltration buffer (10 mM MgCl_2_, 10 mM MES at pH 5.6, 0.1 mM acetosyringone) to the final OD600=0.8. Agrobacterium carrying the constructs of interest were infiltrated in the abaxial side of 4- to 6-week-old *Nicotiana benthamiana* leaves. For cell death phenotype observations, pictures of *N. benthamiana* leaves were taken daily after infiltration. To determine the subcellular localization of the GFP tagged proteins, the leaves were harvested 2 days after infiltration and observed under a Leica TCS SP8 SMD confocal laser-scanning microscope (Ex: 488nm, Em: 500-550nm). Leaves were also collected 2 days after infiltration of two *EIN3* constructs and pGreenII 62-SK as control for quantitative analysis of gene expression.

### Ethylene and ROS treatment

To determine the affects of ethylene on development of shoot tips and pith cavity, 20 mM 1-aminocyclopropanecarboxylic acid (ACC) (Aladdin, Cat No./ID: A101238) and 10 μM aminoethoxyvinylglycine (AVG) (Aladdin, Cat No./ID: A131519) were applied into the culture medium. Shoot tips approximately 1.5 cm in length from water spinach were placed in square culture dishes containing MS medium, MS medium with ACC and MS medium with AVG, respectively. The development of the shoot tips was monitored daily using an X-ray machine (Faxitron UltraFocus60 X).

To investigate whether ethylene and reactive oxygen species (ROS) induce the expression of PCD-regulating genes, 50 µm ACC, 200 mM hydrogen peroxide (H_2_O_2_), and 50 µm methyl viologen (MV) (Aladdin, Cat No./ID: A101238) were sprayed onto the leaves of water spinach. Leaf samples were collected at 3 hours and 24 hours post-treatment for quantitative analysis of gene expression.

### RT-qPCR analysis

The leaves of water spinach and tobacco were ground to a fine powder in liquid nitrogen. Total RNA was extracted using the RNAprep Pure Plant Plus Kit (Polysaccharides & Polyphenolics-rich) (TIANGEN, Beijing, China). The first-strand cDNA was synthesized from 1 μg of total RNA using BeyoRT™ First Strand cDNA Synthesis Kit (RNase H minus) (Beyotime, Shanghai, China). RT-qPCR analysis was conducted with synthesized cDNA and gene-specific primers mixed with the Hieff^®^ qPCR SYBR Green Master Mix (High Rox Plus) (YEASEN, Shanghai, China). To normalize the gene expression, the *Actin* gene was parallel amplified as an internal control. RT-qPCR was performed by ABI StepOne or StepOnePlus Real-Time PCR System (Applied Biosystems, Carlsbad, California, USA). The amplification was performed as follows: 1 cycle of 95°C for 30 sec for initial denaturation, followed by 40 cycles of 95°C for 5 sec, and 60°C for 30 sec. Gene-specific primers used for RT-qPCR are provided in Table S13.

## Data availability

The bulk RNA and single-cell RNA sequencing data are deposited in BIGD under accession number PRJCA031191 (https://ngdc.cncb.ac.cn/gsa/s/1sz3o4XW). The assembled genome have been deposited in the Genome Warehouse in the National Genomics Data Center under accession number PRJCA031243 (https://ngdc.cncb.ac.cn/gwh/Assembly/reviewersPage/LJbeejgROWyPWQrqyFGefceGTtwMdMdKaDMBmTPkArbraimojqSQAhTqJtDSoDRb).

## Acknowledgements

This work was funded by the National Natural Science Foundation of China (32300207 to M.Y., 32472220 to H.W.), the Ministry of Science and Technology of China (2019YFD1000703 to J.Y.), the Shanghai Municipal Afforestation & City Appearance and Environmental Sanitation Administration (G222413 to M.Y., G242407 to W.F., and G222411 to H.W.) and The Science and Technology Commission of Shanghai Municipality (22JC1401300 to H. W.). The authors thank Mr. Binjie Ge from Eastern China Conservation Centre for Wild Endangered Plant Resources (Shanghai Chenshan Botanical Garden) and Mr. Jianhua Ye for providing morphological photos of water spinach. Special thanks to Dr. Lanyun Miao from the Nanjing Institute of Geology and Palaeontology, Chinese Academy of Sciences, for assisting with the illustration. We thank the CEMPS Cell Biology Core Facility in Chenshan site for assistance with Confocal Laser-scanning Microscopy.

## Author contributions

M.Y. contributed to project design, conducted transcriptome and single-cell transcriptome data analysis, performed plant anatomy, histochemical staining, transient tobacco transformation experiments, and wrote the manuscript. F.J. conducted transient tobacco transformation experiments and qPCR analysis. W.Y. constructed the vectors and conducted transient tobacco transformation experiments. J.Z. participated in genome assembly and collecting tissue-specific genes. Y.M. performed paraffin sectioning and TUNEL staining. W.Z. contributed to the transient tobacco transformation experiments. H.Z. performed subcellular localization imaging. Z.X. conducted ethylene reagent treatment, tissue culture, and sample collection. Y.Q.W. contributed to TUNEL staining and collecting tissue-specific genes. Q.H. revised the manuscript. L.Y. provided valuable suggestions and revised the manuscript. H.W. managed the project. J.Y. conceived the project, contributed to genome assembly, and revised the manuscript.

## Competing interests

The authors declare no competing financial interests.

## Notes

### Competing Interest Statement

The authors have declared no competing interest.

